# The timecourse of inter-object contextual facilitation

**DOI:** 10.1101/2023.05.30.542965

**Authors:** Genevieve L. Quek, Alexandra Theodorou, Marius V. Peelen

## Abstract

High-level vision is frequently studied at the level of either individual objects or whole scenes. An intermediate level of visual organisation that has received less attention is the “object constellation” – a familiar configuration of contextually-associated objects (e.g., *plate + spoon*). Recent behavioural studies have shown that information from multiple objects can be integrated to support observers’ high-level understanding of a “scene” and its constituent objects. Here we used EEG in human participants (both sexes) to test when the visual system integrates information across objects to support recognition, using representations of objects’ real-world size as a proxy for recognition. We briefly presented masked object constellations consisting of object silhouettes of either large (e.g., *chair + table*) or small (e.g., *plate + spoon*) real-world size, while independently varying retinal size. As a control, observers also viewed each silhouette in isolation. If object context facilitates object recognition, real-world size should be inferred more effectively when the objects appear in their contextually-associated pairs than in isolation, leading to the emergence of real-world size information in multivariate EEG patterns. Representational similarity analysis revealed that neural activity patterns captured information about the real-world size of object constellations from ∼200 ms after stimulus onset. This representation was stronger for, and specific to, object pairs as compared to single objects, and remained significant after regressing out visual similarity models derived from computational models. These results provide evidence for inter-object facilitation of visual processing, leading to a qualitatively different high-level representation of object pairs than single objects.

**Significance Statement:** This study used electroencephalography decoding to reveal the neural timecourse of inter-object facilitation present for contextually-associated groups of objects (e.g., *chair + table*). Although ubiquitous in daily life, the ’object constellation’ level of representation has rarely been examined compared to isolated objects or entire scenes. By shedding new light on facilitatory interactions between objects, arising before 200ms of visual processing, our results provide insight into the continuum along which objects and scenes exist. At the same time, this work advances the current understanding of the neural basis of real-world size, using strict visual controls to show that inferred information about objects’ spatial scale in the real world emerges around 200 ms after stimulus onset.

## Introduction

Many scenes in daily life are meaningfully defined by the specific constellations of objects they contain (e.g., a knife and fork either side of a plate, or a sofa facing a television). Our encounters with such multi-object arrangements are so ubiquitous that the visual system is remarkably sensitive to statistical regularities within these groups, with both the likelihood of objects’ co-occurrence and their configural arrangement influencing the way we search for, attend to, and remember familiar objects (Biederman, 1972; Biederman et al., 1973; Kaiser et al., 2015; Võ et al., 2019; Nah and Geng, 2022).

Notably, higher-level statistical regularities between objects also appear to influence object processing itself (Kaiser et al., 2019). Behavioural studies have shown that objects are recognized faster and more accurately when viewed together with one or more contextually-associated objects (Auckland et al., 2007; Davenport, 2007). Furthermore, ambiguous objects that appear with an associated object in a familiar configuration (e.g., a keyboard in front of a monitor) are recognized more accurately than the same objects shown in isolation or in an unfamiliar configuration (Bar and Ullman, 1996; Green and Hummel, 2006). In the present study, we use time-sensitive decoding of electroencephalography (EEG) data to examine the timecourse of this inter-object facilitation effect.

Previous EEG studies investigating the timecourse of contextual associations between objects, or between objects and scenes, measured the neural response to semantic violations by contrasting incongruent vs. congruent object associations. This contrast gives rise to reliable event-related potentials (ERPs; Ganis and Kutas, 2003; Mudrik et al., 2010; Võ and Wolfe, 2013; Coco et al., 2020; Quek and Peelen, 2020), including the domain-general N400 over centro-parietal electrodes (Kutas and Federmeier, 2011). Notably, these studies used clearly visible objects, as the interest was in measuring when neural signals reflected the violation. In contrast, here we were interested in how the processing of *ambiguous* objects is facilitated by object context.

A similar question has been addressed for objects disambiguated by global scene context (Brandman and Peelen, 2017; Wischnewski and Peelen, 2021). In those studies, an object was degraded such that it was difficult to recognize when viewed in isolation, but readily identifiable in the context of its original scene. Multivariate decoding analysis showed that the visual representation of the object’s category (animate or inanimate) was facilitated by scene context from ∼300 ms after stimulus onset. Subsequent fMRI and TMS studies revealed that this facilitation resulted from an interaction between processing in scene-selective and object-selective cortex (Brandman and Peelen, 2017; Wischnewski and Peelen, 2021). One possibility is that inter-object facilitation follows the same timecourse as scene-object facilitation, for example because both scene-object and object-object facilitation result from the same feedback mechanism based on the activation of a common conceptual “schema” (Bar, 2004). Alternatively, however, we may expect inter-object facilitation to be faster than scene-object facilitation, as interactions are between representations within the same object processing pathway rather than between separate object and scene processing pathways (Peelen et al., 2023).

To reveal the timecourse of inter-object contextual facilitation, we took advantage of an object property that is rapidly and automatically retrieved following the recognition of objects – real-world size (Konkle and Oliva, 2012b; Hagen et al., 2023). Observers in behavioural and EEG experiments viewed contextually-associated pairs of both small, manipulable objects (e.g., *spoon + plate*) and large, non-manipulable objects (e.g., *desk + chair*). These categories evoke distinct responses in the visual cortex, with large objects activating scene-selective cortex (Konkle and Oliva, 2012a; Coutanche and Thompson-Schill, 2019). However, in addition to reflecting a physical size continuum, this distinction has also been related to objects’ functional properties, reflecting a combination of size and portability (Mullally and Maguire, 2011). For example, large-stable objects evoke a sense of space, while small-manipulable objects do not. Conversely, small-manipulable objects afford grasping and manual actions that large-stable objects do not. Although interesting, exploring how these various conceptual/functional features contribute to the neural distinction between objects of small and large real-world size was not the focus of the present paper. Instead, since neural responses to small and large real-world sized objects are readily decodable using M/EEG (Cichy et al., 2017; Khaligh-Razavi et al., 2018; Wang et al., 2022), we used the presence/strength of real-world size information as a proxy through which to examine how an object’s processing is facilitated by its context. Our rationale was as follows: If object context facilitates object recognition, information about the objects’ real-world size (reflecting physical size/portability/affordances and more) should be inferred more effectively when they appear in contextually-associated pairs than in isolation. To this end, we examined both explicit judgements of objects’ real-world size (Experiments 1a & 1b) and the presence of real-world size information in multivariate EEG patterns (Experiment 2) evoked by viewing objects in silhouette against phase-scrambled noise.

## Materials & Methods

All experimental procedures were carried out with the approval of the Radboud University Faculty of Social Sciences Ethics Committee (ECSW2017–2306-517).

### Participants

For Experiment 1a (behavioural), we recruited 75 online participants (40 female, 33 male, 2 not answered) aged between 18-35 years (*M* = 26.31, *SD* = 4.89) via Prolific (www.prolific.co). All were living in the Netherlands, had normal or corrected-to-normal vision, had prior experience in online research participation (>10 studies), and were monetarily compensated for their participation. After applying participant qualification criteria (see Analysis), the final sample for Experiment 1a was 64. For Experiment 1b (behavioural), 91 undergraduate students from the University of Western Sydney (78 female, 12 male, 1 not answered) aged between 17-45 years (*M* = 21.2, *SD* = 5.76) participated in the online experiment in exchange for course credit. The final sample after applying participant qualification criteria was 60. For Experiment 2 (EEG), we used G*Power 3.1.9.2 (Faul et al., 2009) to compute the sample size required to give 80% power to detect a medium effect size (Cohen’s d = 0.5) in a one-sample test (α=0.05, two tailed). We used a one-sample test for this power analysis since we ultimately aimed to evaluate whether beta coefficients given by multiple regression were significantly greater than zero. This indicated a required final sample of N=34, which we achieved (after participant exclusion) by testing 41 English-speaking individuals (11 males) aged between 18 and 35 years (*M* = 23.46, *SD* = 3.88) from the local region of Nijmegen, the Netherlands. All had normal or corrected-to-normal vision, with no history of neurological illness/concussion/brain surgery/migraine. Data from seven participants were discarded to reach the eventual N=34: this included one participant who terminated the experiment early due to discomfort, and six participants with very high noise/muscle artefact in the signal.

### Stimuli

Stimuli were identical across the two experiments and were based on 20 pairs of contextually-associated objects, each denoting either a small- or large-scale object constellation (Figure 1A). ‘Small’ displays contained pairs of desktop-sized objects that could fit into a shoebox, typically with grasp/reach affordances (e.g., *bottle + glass*). ‘Large’ displays contained pairs of furniture-sized objects, typically with navigational affordances (e.g., *table + chair*). All objects appeared in silhouette, embedded in a noise mask, with no other meaningful visual information (e.g., background). These sparse visual displays served to eliminate many of the inherent low- and mid-level visual differences between naturalistic scenes of different spatial scales. Note that such visual controls do not eliminate more conceptual differences between desktop-size and furniture- size objects – for example, their respective manipulation and navigation affordances. Importantly, where these and other distinctions between large and small objects could be problematic for isolating neural signals corresponding to real-world size *per se,* they do not impede our ability to treat real-world size judgements/representations as a useful index of object processing and recognition.

**Figure 1.**
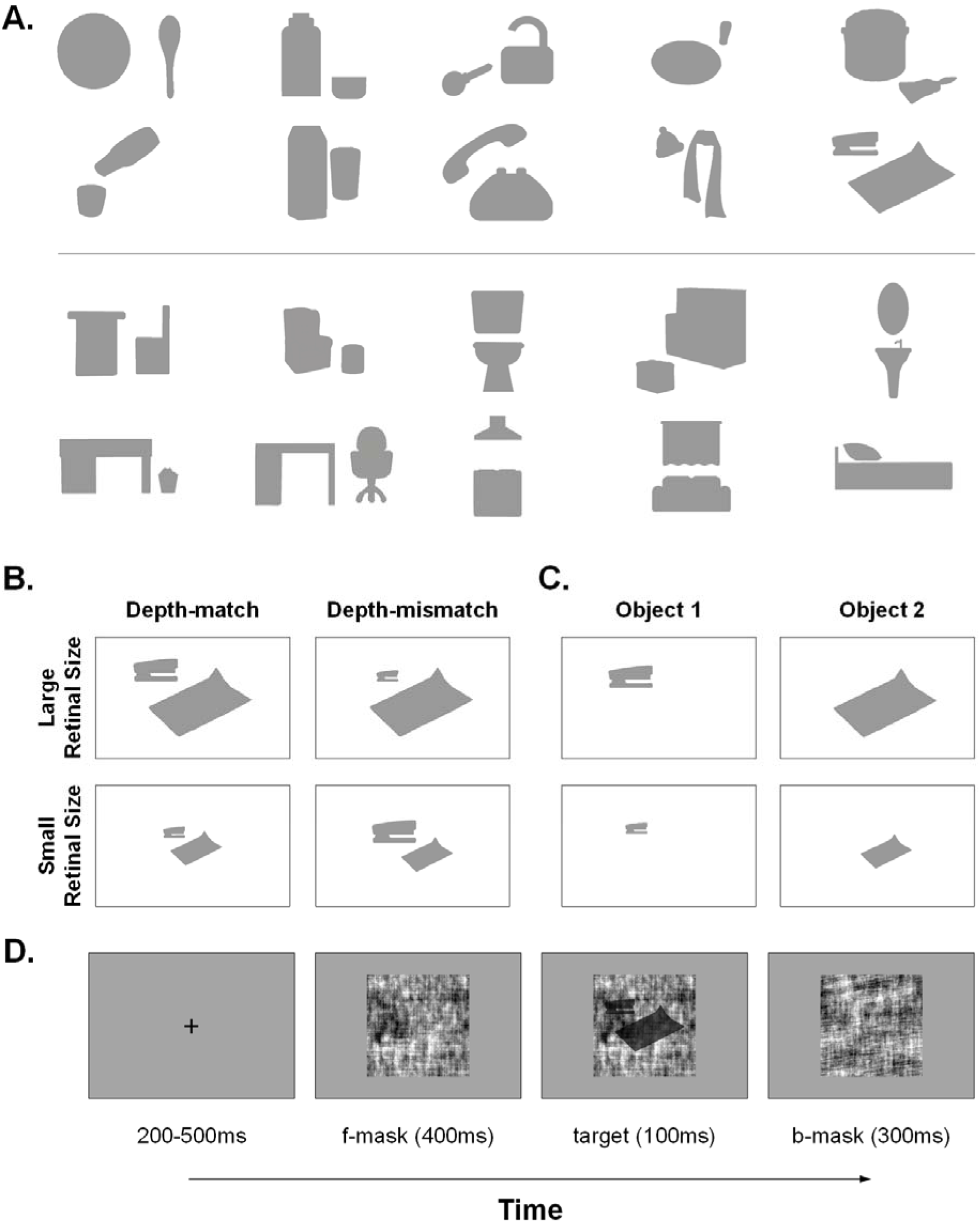
Stimuli. A) The 20 base object pairs drawn from small (top row) and large (bottom row) real-world contexts. B) Each object pair had four versions: A large and small retinal size in which the objects’ relative sizing implied they were at the same depth (Depth-match). Recombining the elements of these versions produced two pairs where the objects were implied to be at different depths (Depth-mismatch). Retinal size labelling for these Depth-mismatch pairs following the retinal size of the more dominant object in the pair (see Methods). C) Participants also saw each individual object in isolation. D) Basic trial structure for all three experiments. Following fixation, the target appeared at 40% contrast embedded in a phase-scrambled noise mask, and was immediately followed by a different phase-scrambled mask. In Experiments 1a & 1b, the fixation period was fixed at 500 ms.

Inspired by previous work showing that relative spatial positioning modulates the extent to which observers group/integrate concurrently presented objects (Bar and Ullman, 1996; Baeck et al., 2013; Kaiser et al., 2019; Xu et al., 2021), we varied the relative sizing of the objects within each pair as depicted in Figure 1B. We began with the stimuli shown in Figure 1A, wherein the proportional relative sizing of the object silhouettes within each pair implied them to be at the same distance or depth. We then created an identical ‘depth-match’ version of the pair at a decreased retinal size (2:1 size decrease), and recombined the elements of these two versions to create two different ‘depth-mismatch’ pairs in which the objects’ disproportionate retinal sizes (e.g., large *pillow* + small *bed*) implied they were at different distances from the viewer. To ensure that the pixel distance between the two objects in a depth-mismatch pair was equivalent to the pixel distance between their depth-match counterparts, we randomly labelled the objects in each pair as object 1 and object 2. We then utilized the smaller distance whenever object 1 was the small object in the depth-mismatch pair and the large distance when it was the larger object. We maintained the surface alignment of the two-object scene as much as possible across versions, e.g., since *desk* and *chair* both rest on the floor, we aligned these items at their bottom edge regardless of whether their depths were matched or mismatched. Large/Small retinal size labels for the depth-mismatch pairs followed the retinal size category of the larger, more dominant object within the original pair. For example, depth-mismatch versions of the *paper + stapler* pair were labelled according to the retinal size of *paper*, since this was the more dominant object in the original, depth-match pair (see Figure 1B). Thus, while depth-mismatch pairs were inherently comprised of one small version object and one large version object, mismatch pairs labelled as small retinal size had significantly fewer pixels (*M* = 26368 pixels, *SD* = 9500 pixels) than mismatch pairs labelled as large retinal size (*M* = 46109 pixels, *SD* = 15476 pixels), *t*(19) = 6.32, *p* < .00001.

In addition to the four versions of each object pair, observers also saw the small and large retinal size versions of every silhouetted object in isolation (Figure 1C). The position of each isolated object at a given retinal size was always halfway between where that object had appeared as part of the depth-match and depth-mismatch pair. This made for a total of 80 object pairs and 80 single objects (10 exemplars x 4 versions representing each level of real-world size). There were 248 phase scrambled noise images, two of which were randomly selected to serve as masks on each trial.

Trial structure was near identical across Experiments 1 and 2 (Figure 1D): The trial began with a central fixation cross (duration of 500 ms in Experiment 1; between 200-500 ms in Experiment 2). Following the fixation period, a phase-scrambled noise mask appeared for 400 ms, then the silhouette target appeared at 40% contrast embedded within the mask for 100 ms. The trial concluded with a different phase-scrambled backward mask that lasted 300 ms. In both experiments, the trial sequence appeared against a uniform grey background.

### Design & Procedure

In Experiment 1a (behavioural), we were interested in participants’ ability to explicitly categorise the real-world size of object silhouettes presented in either contextually-associated pairs or as singletons. Our rationale was that the objects would be more readily recognised when they appeared in familiar constellations than when they appeared alone – and that this should result in faster/more accurate categorisation of their real-world size. The experiment was programmed in PsychoPy (jsPsych) and hosted online on Pavlovia (Open Science Tools Limited, 2021). Participants completed the experiment in a browser window on either a desktop or laptop computer. Before beginning, we instructed them to minimise any distractions in the environment (e.g., music, TV, other people), and to disconnect any external monitors to ensure the experiment appeared on their main screen. They were told that they would see briefly presented object silhouettes, and that the task was to decide how large those object(s) would be in the real world. On each trial (see Figure 1D), participants had 1000 ms from target onset to categorise the displayed object(s) as either large or small by pressing either the *f* (small) or *j* (large) key (response key labels appeared onscreen throughout the trial). Importantly, the task instructions included an explicit reference object to help participants delineate the boundary between objects of large and small real-world size: participants saw three example silhouette stimuli representing each real-world size, annotated with the labels “The objects could be relatively small, i.e., could fit in a shoe box” and “The objects could be relatively large, i.e., could not fit in a shoebox”, and could review these instructions for as long as they wished before pressing SPACE to continue. We also specifically instructed participants to focus on the size of the objects in the <u>real world</u> (not the size at which they appeared onscreen), and to fixate on the central cross whenever it appeared between trials. There were 16 practice trials where the target duration was set at 500 ms to facilitate learning; these training trials used an independent stimulus set to the main experiment. In the experiment proper, 80 object pairs and 80 single objects appeared in random, intermingled order for a total of 160 trials. Mask stimuli were sized at 560×560 pixels; we could not fully control the visual angle subtended by the stimuli owing to conducting the study online. Total testing time for Experiment 1a was approximately 10 minutes including a self-paced break at the halfway point.

Experiment 1b (behavioural) was identical to Experiment 1a in all respects, save that here half the trials comprised the large retinal size versions of the original object pairs (20 stimuli), while the other half comprised random pairings of objects drawn from the same real-world size category, but from different scene contexts (e.g., *bed + stove*). Each object pair was presented once for a total of 40 trials; the total testing time was approximately 5 minutes.

Experiment 2 (EEG) was conducted in the lab. The trial sequence was programmed in Presentation® (Neurobehavioral Systems, Inc., Berkeley, CA, www.neurobs.com), and displayed on a BenQ XL2420Z computer monitor (120Hz refresh rate; 1920×1080 resolution). Mask stimuli were sized 14×14cm, with viewing distance set at 65cm such that each stimulus subtended 12.3 x 12.3 degrees of visual angle. Experiment 2 saw a 10-fold increase in the number of trials: each of the 160 unique stimuli appeared once per block, with 10 blocks in the full experiment. In contrast to Experiment 1, here participants were engaged in a 1-back task, monitoring the target from trial to trial and pressing the spacebar whenever they saw an exact image repeat (10 instances per block). Prior to the main experiment, observers performed 25 practice trials with a slightly longer target duration (300 ms); this included 5 repeat instances to enable them to learn the task. The full experimental session including EEG set up lasted around 2.5 hours.

We recorded scalp EEG in Experiment 2 using a 64-channel BrainProducts actiCAP active electrode system with a sampling rate of 500Hz (Brain Products GmbH, Gilching, Germany). We used customized electrode positions adapted from the actiCAP 64Ch Standard-2 system (ground placed at AFz; TP10 placed on right mastoid). Impedances for individual scalp channels were held <40 kOhm, with data referenced online to the left mastoid and filtered between 0.016 and 125 Hz using BrainVision Recorder (Brain Products GmbH, Gilching, Germany). We monitored participants’ eye movements via external passive electrodes situated at the outer canthus of each eye, as well as immediately above and below the right eye. The ground electrode for passive ocular channels was placed on the tip of the nose. The experimenter visually monitored the EEG trace throughout the experiment, initiating each experimental block manually after ensuring there were no high amplitude deflections resulting from ocular/muscular artefacts. Stimulus onsets were marked in the EEG file using numeric triggers.

### Analysis

#### Experiment 1a & 1b

For the behavioural real-world size categorisation task, we reached a final sample of 64 after removing participants who failed to provide a response on more than 30% of trials (*n* = 5), and subsequently any participant whose mean accuracy or response time (RT) on correct trials fell more than 2 standard deviations below the group mean (*n* = 6). We further discarded any trials on which the participant responded sooner than 200 ms following target onset (0.62% of trials), under the rationale that these early responses were unlikely to be informative. We then inspected *i)* response accuracy (treating trials with no response as incorrect responses) and *ii)* RT on correct trials. For the latter, we bounded each participant’s RT distribution at 2 standard deviations above and below the mean by replacing values outside this range with the top and bottom RTs respectively (4.6% of trials). For each metric, we ran a 2 x 2 repeated measures ANOVA with the factors *Stimulus Type* (Pair, Single) and *Retinal Size* (Large, Small). Note that for the depth-mismatch pair stimuli, the ‘large’ and ‘small’ retinal size labels were simply arbitrary labels, since each depth-mismatch pair necessarily comprised one large retinal size object and one small retinal size object (see Figure 1B). A subsequent analysis inspected the effect of *Relative Depth* (match, mismatch) for the Object Pairs data alone (since there was no relative depth manipulation for single objects).

Analysis for Experiment 1b followed this model closely: the same exclusion criteria resulted in a final N of 60 participants (27 removed due to >30% of trials without a response; 4 removed as outliers). After trial trimming, we compared responses to matched object pairings and random object pairings using a one-tailed paired samples *t*-test for *i)* response accuracy (treating trials with no response as incorrect responses) and *ii)* RT on correct trials.

#### Experiment 2

##### EEG pre-processing

We pre-processed the EEG data in MATLAB 2016b using FieldTrip functions (Oostenveld, Fries, Maris & Schoffelen, 2011; http://fieldtriptoolbox.org). This included a bandpass filter between 0.05-100 Hz, a line noise filter at 50 Hz, 100 Hz, & 150 Hz, and downsampling the data to 250 Hz for easier storage and handling. A maximum of three artefact ridden channels were identified by eye and interpolated using the weighted average of neighbouring electrodes. To combat artefacts introduced by eyeblinks, we relied on an independent components analysis approach (runica decomposition), visually inspecting the components and associated topographies for each participant to identify and remove the eye blink component. No trial or epoch rejection was applied. We then re-referenced each participant’s cleaned EEG trace to the average of all channels before segmenting an epoch from -200 ms to 500 ms around the onset of each target stimulus (excluding 1-back repeat instances). The resulting 800 single object epochs and 800 object pair epochs were then baseline-corrected using the 200 ms prior to stimulus onset.

##### Representational Similarity Analysis (RSA)

###### RSA using individual versions of each object pair

For the object pair neural data, we used RSA (Kriegeskorte et al., 2008; Kriegeskorte and Kievit, 2013) to build a pairwise representational dissimilarity matrix (RDM) at each 4 ms timepoint, for each participant separately. This involved pairwise decoding of every version of every object pair exemplar (i.e., 80 items) using Linear Discriminant Analysis (LDA) with 10 fold cross-validation (leave-one-block-out) as implemented in the CoSMoMVPA toolbox (Oosterhof et al., 2016; https://www.cosmomvpa.org/). We used multiple regression to relate the resulting time-varying RDM to four categorical predictors corresponding to the binary models given by our experimental design (see Figure 3B). These were <u>Pair Identity</u> (i.e., the specific real-world context the objects were drawn from), <u>Retinal Size</u> (i.e., large or small version), <u>Real-World Size</u> (i.e., small- or large-scale objects), and <u>Relative Depth</u> (i.e., implied depth of the objects either matched or mismatched within the pair). Here and in all following analyses, we inspected the utility of each model as a predictor of the neural RDM at each timepoint between 0 and 500 ms via a Bayesian one sample *t*-test that examined whether the participant mean betas were greater than zero (Teichmann et al., 2022). The resulting Bayes Factors (BFs) centre around 1, with BF<1 comprising evidence for H₀ (mean beta equal to zero) and BF>1 comprising evidence for H_1_. Importantly, BFs can be directly interpreted as how much more likely the observed data are under this alternative hypothesis than the null (Morey et al., 2016). Thus, the larger the BF, the stronger the evidence for H_1._ Here we chose to depict timepoints where there is moderate evidence one way or the other, i.e., BF ≤ 1/3 (positive evidence for H_0_) or BF ≥ 3 (positive evidence for H_1_) (Andraszewicz et al., 2015). No further correction for multiple comparisons was implemented (see Teichmann et al., 2022).

###### RSA using object pairs collapsed across version

Next, we ran a simplified RSA that pooled the neural data for the four versions of each object pair (see Figure 1B) for pairwise decoding. This effectively collapsed across all design factors other than Real-World Size, which remained the primary model of interest. Here we aimed to inspect the predictive power of the real-world size model while controlling for the inherent low-level differences between large and small object pairs (Robinson et al., 2023). To this end, we built four low-level visual control RDMs (see details below), each sized 80×80 (i.e., every version of every object pair treated individually). We then collapsed these into the reduced 20×20 RDMs by averaging values over the four versions of each object pair, e.g., collapsed dissimilarity value for Pair 1 vs. Pair 2 was the average of all RDM intersections of [1A, 1B, 1C, 1D] and [2A, 2B, 2C, 2D]. The visual control models we considered were: *1)* An image similarity model reflecting 1 minus the pixel-wise correlation between each of the 80 object pair images; *2)* A silhouette overlap model (Jaccard, 1901) comprising 1 minus the Jaccard similarity (ranges between 0 and 1) for each pairwise combination of the 80 object pair images; 3) A computational model of low-level visual processes, obtained by applying HMAX (Riesenhuber and Poggio, 1999; Serre et al., 2007; https://maxlab.neuro.georgetown.edu/hmax.html) to each of the 80 object pair images embedded in an example noise mask (just as they appeared in both experiments). RDM values were 1 minus the Spearman correlation between the vectorised responses on the final HMAX C2 layer; and 4) A mid-level feature model comprising the pairwise Euclidean distances between curvature indices for object pairs obtained using a computational model developed by Li and Bonner (2020; https://github.com/shipui2005/Curvature-Model). In this and the following multiple regressions, we inspected the Spearman rank-order correlations between all model RDMs, applying a significance criterion of *p <*.01, FDR corrected. Since the <u>Pixel Correlation</u> and <u>Jaccard Similarity</u> models were highly correlated (rho = 0.9844, *p <*.0001) and displayed large Variance Inflation Factors (VIFs) (pix = 31.09, jac = 39.91), we elected to drop the <u>Jaccard</u> model from our multiple regression, leaving 1 categorical predictor (<u>Real-world Size</u>) and 3 continuous control predictors (<u>pixel correlation</u>, <u>HMAX C2</u>, and <u>curvature</u>, see Figure 4). Since the intercorrelations between these predictors were not especially high (only <u>Pixel Correlation</u> and <u>Curvature</u> were significantly correlated, rho = 0.2587, *p* = .002), and since VIFs were all close to 1 (RwS = 1.0046, pix = 1.0645, hmx = 1.0262, cur = 1.0884), we felt justified in including these models as independent predictors in the same multiple regression.

### Comparisons with Single Objects

If recognising an object is facilitated by its inclusion in a familiar object constellation, then the representation of real-world size it evokes will be stronger than that evoked by the same object viewed in isolation. To enable this comparison, we duplicated the above collapsed RSA pipeline for the Single Object neural data, producing corresponding real-world size and low-level visual control models for the single object stimuli. For the singles objects, we found significant correlations between the real-world size and curvature models (rho = 0.2516, p = .0028) and the curvature and pixel correlation models (rho = 0.2276, p = .0049). Importantly, VIFs for the predictors were all close to 1 (RwS = 1.0884, pix = 1.0885, hmx = 1.0251, cur = 1.1608), such that we felt confident interpreting the beta estimates from a multiple regression that included these models as independent variables.

### Comparisons with a Deep Convolutional Neural Network

To further rule out the possibility that the neural distinction between large and small real-world size object pairs arises due to low-level featural differences between large and small objects, we examined representations of these sparse scenes in the layers of a deep convolutional neural network trained to label objects using 1.2 million naturalistic images (AlexNet, Krizhevsky et al., 2012). We obtained output activations corresponding to the 80 object pair images embedded in an example noise mask (i.e., as they appeared to human observers in the two experiments) on each of the 8 AlexNet layers (input layer, five convolutional layers, two fully connected layers), each of which was used to build an RDM. Just as for the low-level visual control models, we collapsed the resulting 80×80 RDMs down to 20×20 by averaging values over the four versions of each object pair. Using the same significance criterion as detailed above, we inspected the intercorrelation matrix for these eight AlexNet RDMs and the model of Real-world Size. If featural differences between our large/small stimuli account for the predictive power of real-world size, then we would expect strong correlations between this model and the earlier layers of AlexNet.

Lastly, given that the upper layers of convolutional neural networks capture some important organising principles of object vision (e.g., animacy, Khaligh-Razavi and Kriegeskorte, 2014), we also asked whether the information about real-world size in our neural data could be accounted for by the representational structure evident in these layers of AlexNet (i.e., FC6 & FC7). To this end, we subjected the (reduced) neural RDM series for object pairs to separate multiple regressions that included <u>Real-world Size</u> as a categorical predictor alongside i) the FC6 RDM, and 2) the FC7 RDM (treated as continuous predictors). Although the <u>Real-world Size</u> model was significantly correlated with both AlexNet layer models (FC6: rho = 0.20, *p* = 0.0079; FC7: rho = 0.24, *p* = 0.0016, FDR-corrected), Variance Inflation Factors (VIF) for these regressors were close to zero in both cases (RwS & FC6 = 1.0378; RwS & FC7 = 1.0391). As such, we felt justified in treating them as independent predictors in these multiple regressions.

## Results

### Experiment 1a: Behavioural judgements of real-world size

To verify that the object silhouettes were recognised better when presented in pairs than in isolation (Bar and Ullman, 1996), we compared participants’ ability to determine the real-world size of the silhouettes between these conditions. A 2×2 repeated measures ANOVA for conditional mean accuracies confirmed that participants could determine the real-world size of object silhouettes significantly better when the items appeared in contextually-related pairs that formed sparse scenes (*M* = 77.3% correct) rather than in isolation (*M* = 65.32% correct; main effect of *Stimulus Type*, *F*(1,63) = 194.72, *p <*.00001, ges = 0.512, see Figure 2A). This was qualified by a significant interaction with *Retinal Size*, *F*(1,63) = 17.40, *p <*.0001, ges = .084, wherein the magnitude of the Pair > Single difference was larger for the small retinal size versions (*M*_diff_ = .155, *SD* = .096) than the large retinal size versions (*M*_diff_ = .084, *SD* = .098). The main effect of *Retinal Size* was not significant, *F*(1,63) = 0.25, *p* = .619, ges = .001. For RT (Figure 2B), we observed a similar main effect of *Stimulus Type*, *F*(1,63) = 8.16, *p <*.01, ges = .041. On average, participants were slower to categorise the real-world size of single objects (*M* = 595 ms, *SD* = 139 ms) compared to object pairs (*M* = 590 ms, *SD* = 136 ms). Neither the main effect of *Retinal Size, F*(1,63) = 3.49, *p* = .467, ges = .003, nor the two way interaction, *F*(1,63) = 0.53, *p* = .066, ges = .016, reached significance in the case of RT. Together, these results indicate that observers recognised objects (and could therefore access information about their real-world size) better when the objects appeared as part of a familiar constellation than when they appeared alone, replicating previous work (Bar and Ullman, 1996).

**Figure 2.**
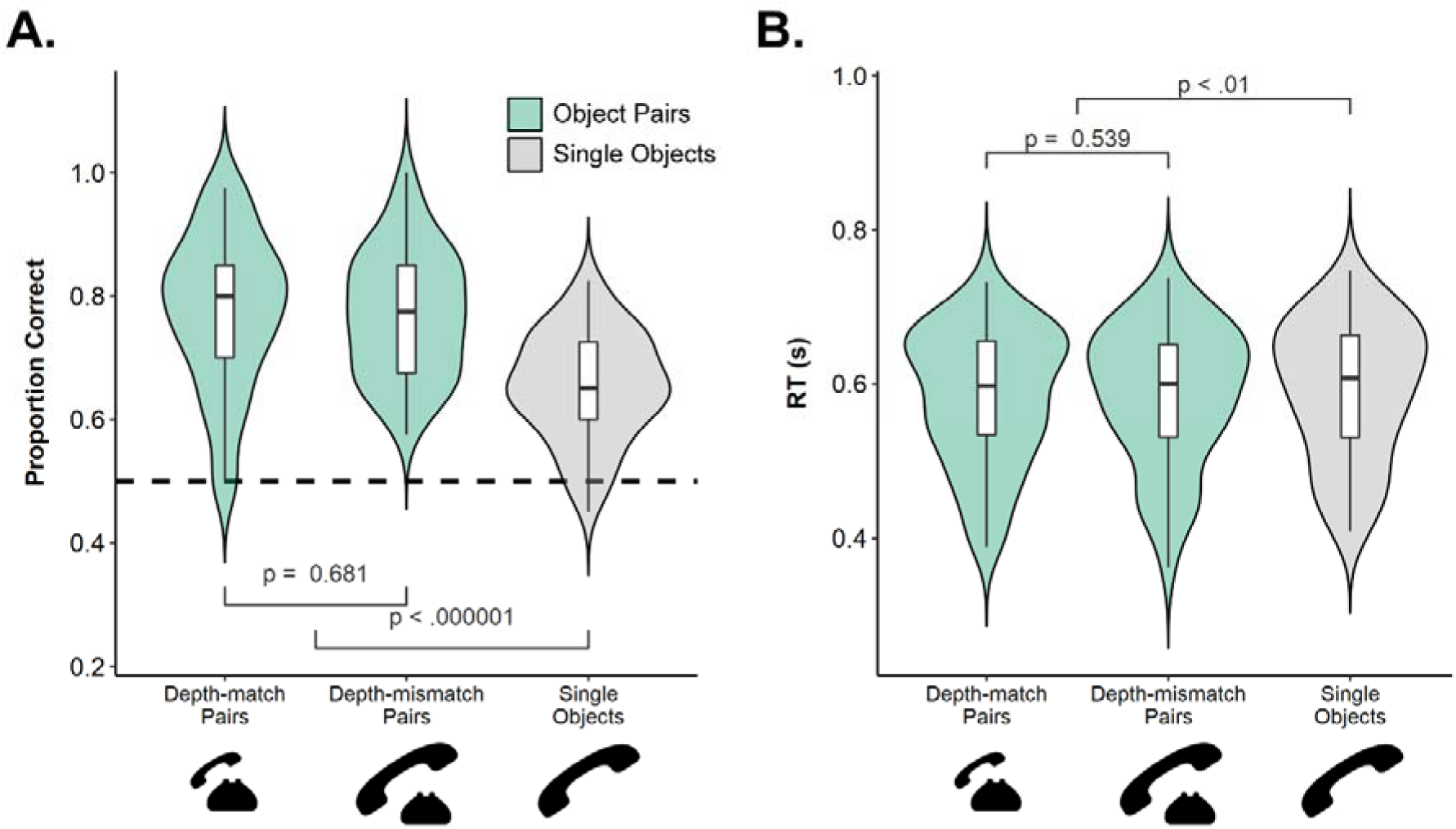
Behavioural judgements of real-world size in Experiment 1a. Overlaid violin and box-and-whiskers plots of A) accuracy and B) RTs. Single Object data (in grey) appear at right, with Object Pair data (in green) split into depth-match and depth-mismatch versions (there was no significant difference between these conditions in either metric). Upper and lower box edges correspond to the first and third quartiles (capturing the interquartile range, IQF); whiskers encompass values less than 1.5*IQF from the box edges. Example stimuli for each condition appear below the x-axis labels.

Contrary to our predictions, there was no evidence in accuracy or RT that participants’ real-world size judgements were affected by the relative size of objects within a pair (Figure 2). Paired *t*-tests of both accuracy, *t*(63) = 0.413, *p* = .681, and RT, *t*(63) = -0.62, *p* = .539, revealed no difference between responses to depth-match and depth-mismatch object pairs. Note that there was no such comparison for single objects, as relative depth does not apply to isolated objects.

### Experiment 1b: Control

One possibility is that the advantage that object pairs enjoy over single objects has nothing to do with pairs comprising familiar object-constellations, but rather simply arises because pairs contain twice as much real-world size information as single objects (i.e., ‘two objects are better than one’). If this were the case, then judgements of real-world size should be comparable for *any* pair of objects representing the same spatial scale, regardless of whether they form a contextually-associated constellation or not.

To assess this, we ran an additional behavioural control experiment using novel small and large object pairs formed by randomly combining objects across scene contexts (e.g., *cup + scarf*; *stove + curtain*). Each of these 20 random object pairs (10 representing each RWS, only large retinal size versions included) and the original 20 contextually-associated object pairs (large retinal size versions) appeared once in an online experiment (randomly intermingled). Experiment and task parameters were otherwise identical to Experiment 1a. A one-tailed paired samples *t*-test indicated that participants did indeed judge the real-world size of object silhouettes more accurately when they appeared as part of a familiar object constellation (*M* = 68.14%) than when they appeared alongside a random object from the same size category (*M* = 63.98%), *t*(59) = 2.33, *p* = .012. There was no evidence that RT differed significantly between the randomly combined pairs (*M* = 589 ms, *SD* = 102 ms) and contextually-associated pairs (*M* = 580 ms, *SD* = 103 ms), *t*(59) = - 1.33, *p* = .094.

### Experiment 2: The timecourse of inter-object facilitation

In Experiment 2, we applied RSA to our EEG data to examine the temporal dynamics of neural representations evoked by large and small real-world size object silhouettes. As above, our primary interest was in understanding whether concurrently-presented objects facilitate each other during visual processing. Here we reasoned that if presenting objects in familiar constellations facilitates object processing, then the neural response to object pairs should carry information about their associated real-world size to a greater extent than the response to single objects.

#### RSA for the individual versions of each object pair

We began by inspecting how well the binary models based on our design factors (Figure 3B) could predict the neural RDM series for object pairs (Figure 3A). In this analysis, we treated the four versions of each object pair (e.g., stapler + paper) independently, building an 80×80 RDM for each timepoint (see Methods for details). Multiple regression beta estimates over time for each of the four predictor models are given in Figure 3C, together with Bayesian one-sample t-tests evaluating beta > 0.

**Figure 3.**
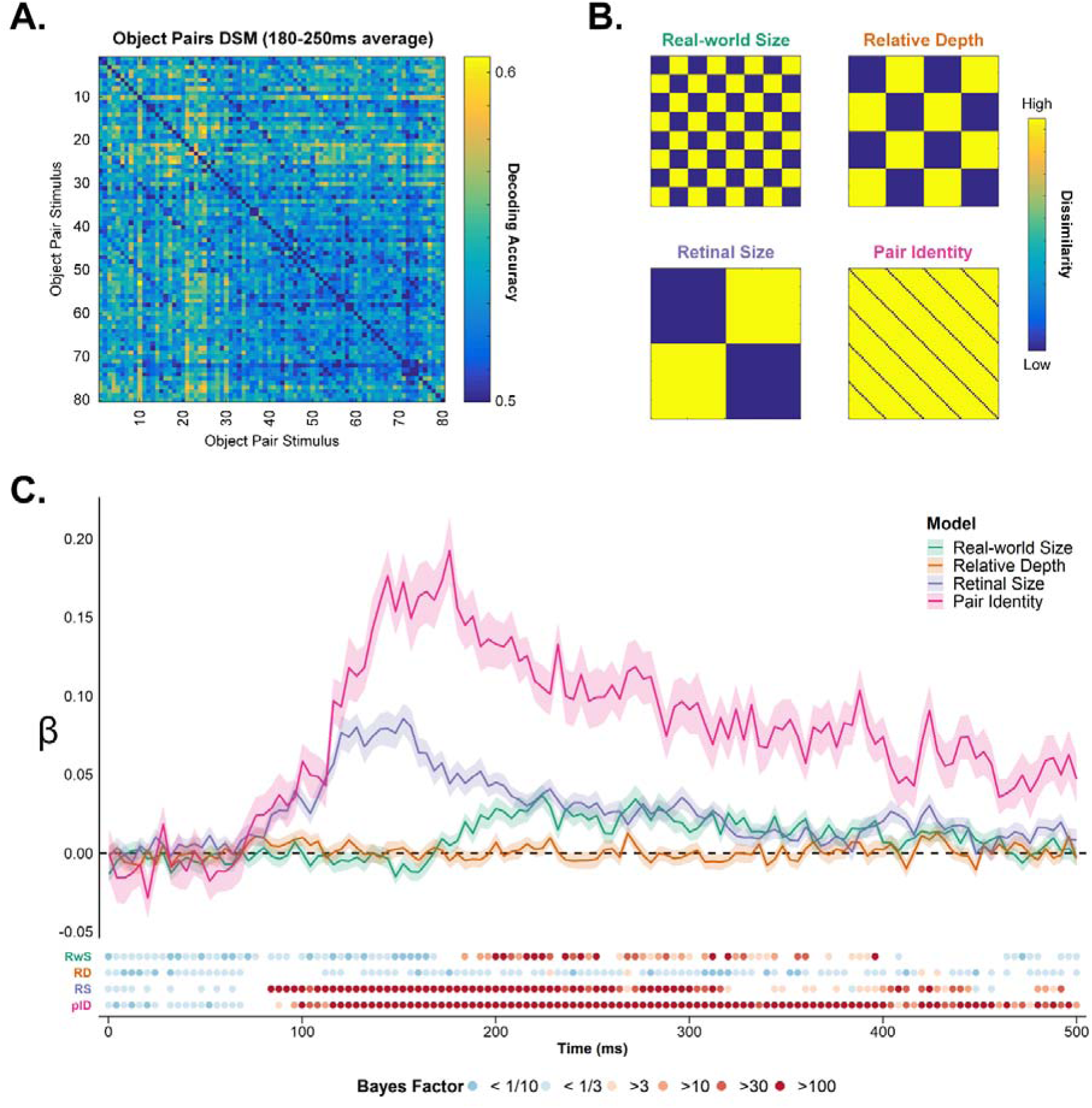
RSA for 80 individual object pair stimuli in Experiment 2 (i.e., treating each of the 4 versions for the 20 object pairs separately). **A)** The neural RDM averaged over the epoch between 180-250 ms. **B)** Binary RDMs for the stimulus design factors treated as categorical predictors in a multiple regression of the neural RDM series. **C)** Beta estimates for the four predictors included in the multiple regression, plotted as a function of time from stimulus onset. Shaded regions are within-subjects standard error. Lower panel shows colour coded Bayes Factors for Bayesian one-sample t-tests comparing beta against zero at each timepoint, for each predictor.

Two of these models predicted the neural RDM from a relatively early stage of stimulus processing: <u>Retinal Size</u> (onset = 84 ms, peak = 152 ms) and <u>Pair Identity</u> (onset = 88 ms, peak = 176 ms). This was expected, since both models capture visual features inherent to the stimuli such as pixel coverage and global shape. Notably, information about both the object identities and their display size was also present in the neural response at much later timepoints, suggesting that the visual system maintains representations of both of these dimensions for some 500 ms after the objects come into view.

In contrast to the <u>Pair Identity</u> and <u>Retinal Size</u> models, the <u>Real-world Size</u> model requires similar neural responses across stimuli with both distinct object identities and visual features (i.e., the response to *hat + scarf* has to resemble that of *stapler + paper*, while also generalising across large and small versions of those pairs). This real-world size model emerged as a relevant predictor of the neural response at a comparatively later stage of stimulus processing (onset = 184 ms, peak = 224 ms), and remained a relevant predictor until as late as 400 ms.

The other high-level model yielded by our design was <u>Relative Depth</u>, which captured whether the objects within a pair were implied to be at the same or different depth from the viewer. As is clear in Figure 3C, this model was not a significant predictor of the neural data at any timepoint, suggesting that the objects’ relative sizing in each display did not modulate the neural response. <u>Relative Depth</u> was also not a relevant predictor of the object pair neural data when considered as an isolated categorical predictor in a separate regression analysis (data not shown). This was consistent with behavioural observations from Experiment 1, where we found no evidence that the relative depth of objects within a pair influenced participants’ real-world size judgements.

#### RSA for object pairs collapsed across version

Since the relative depth of objects within a pair modulated neither the neural response to our object stimuli nor participants’ overt categorisations of real-world size (Experiment 1), we next ran a simplified RSA in which the depth-match and depth-mismatch versions of every object pair were treated as a group.

Collapsing across version in this way had the effect of eliminating the <u>Relative Depth</u>, <u>Retinal Size</u>, and <u>Pair Identity</u> models (the latter becomes the RDM diagonal), leaving only the <u>Real-world Size</u> model. We evaluated the utility of this categorical predictor in a multiple regression analysis alongside three low-level visual control models included as continuous predictors (see Methods). Our rationale here was to isolate information about real-world size carried in the neural signal that could not be explained by inherent differences in visual features between the object constellations representing small and large spatial scales. Since information about objects’ real-world size should become accessible only after the objects are recognised, representations of real-world size that are controlled for visual feature differences serve as a useful proxy for object processing.

Figure 4 shows the results of this multiple regression analysis collapsing over the four versions of each object pair. <u>Real-world Size</u> did not correlate with any of the three low-level visual control models (**pix:** rho = -0.04, *p* = .608; **hmx:** rho = -0.05, p = .608; **cur:** rho = 0.05, p = .608, FDR-corrected). Bayesian one-sample *t*-tests indicated that the model which predicted the neural response earliest in time was <u>Pixel Correlation</u> (onset = 104 ms, peak = 156 ms), followed by the <u>HMAX (C2)</u> (onset = 148 ms, peak = 176 ms). These early onsets of predictive power were consistent with these models capturing perceptual features of the images, which should be relevant to the feed-forward sweep of visual processing. We did not find any evidence that the <u>Curvature</u> model contributed unique predictive power to the neural response, although it was a meaningful predictor when treated as the sole factor in a regression of the neural RDM series analysis (as were all the low-level visual control models; data not shown).

**Figure 4.**
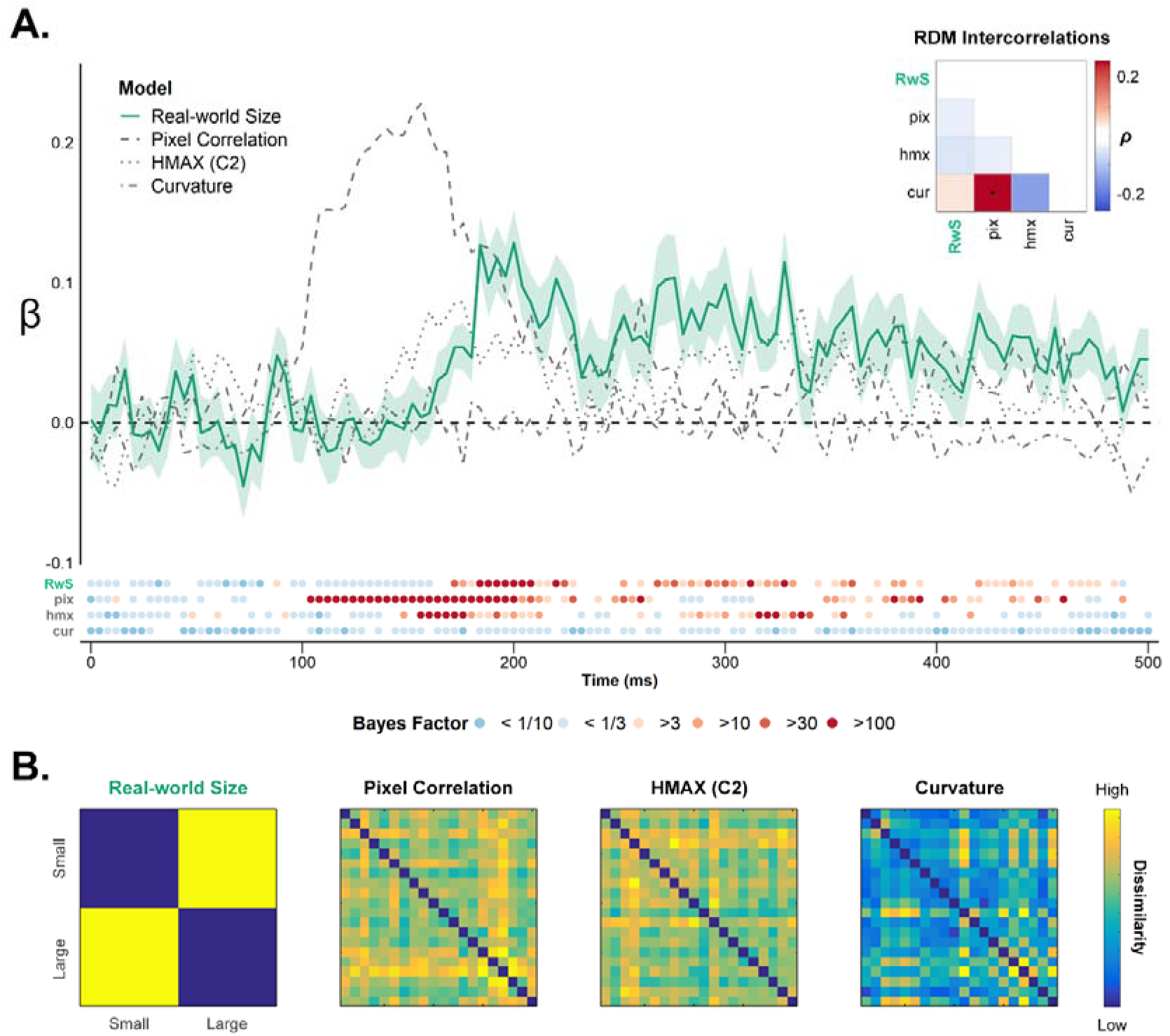
RSA for the 20 object pairs pooled across the four versions (see Figure 1B). A) Multiple regression betas for Real-world Size (in green) and low-level visual control predictors (in grey) of the neural RDM series. Shaded region for the RWS model is within-subjects standard error. Lower panel shows colour coded Bayes Factors for one-tailed *t*-tests (mean beta > 0) at each timepoint, for each predictor. Inset: Model intercorrelation matrix; asterisks indicate Spearman correlations significant at p <.01 (FDR corrected). Real-world Size was not significantly correlated with any of the low-level feature models. B) Models RDMs. Note that all models were confirmed as relevant predictors when treated as an isolated predictor of the neural RDM series (not shown here).

Our primary interest in this analysis was to determine whether the <u>Real-world Size</u> model would offer any unique explanation of the neural response when evaluated in the context of various low-level visual control models. This was indeed the case, with <u>Real-world Size</u> remaining a significant predictor even when simultaneously regressing out three visual control models (Figure 4A). The timecourse was similar to that observed in the previous analysis (onset = 172 ms, peak = 200 ms), and remained intermittently relevant to the neural response throughout the period examined here (i.e., up to 500 ms).

#### Comparison with single objects

In Experiment 1, observers were less able to judge the real-world size of objects when viewed in isolation compared to when they appeared in related pairs. Here we asked whether the neural response to single objects contained information about their real-world size, once again controlling for visual feature differences between large and small objects. We ran the same multiple regression analysis as for Object Pairs, in which <u>Real-world Size</u> was a categorical predictor alongside three low-level visual control predictors derived from the single object stimuli (i.e., <u>pixel correlation</u>, <u>HMAX (C2)</u>, & <u>Curvature</u>). Figure 5 shows the resulting beta- timecourse for the <u>Real-world Size</u> model of Single Object data, with the corresponding beta-timecourse for object pair neural data overlaid in green. In summary, there was no evidence that <u>Real-world Size</u> was a relevant predictor for the single object neural response after controlling for low-level visual features of the objects (although this model was relevant when treated as single predictor in the regression, results not shown here). <u>Real-world Size</u> was also a significantly better predictor for object pairs than for single objects around 200 ms after stimulus onset (Figure 5), providing a possible neural correlate for the behavioural effect observed in Experiment 1.

**Figure 5.**
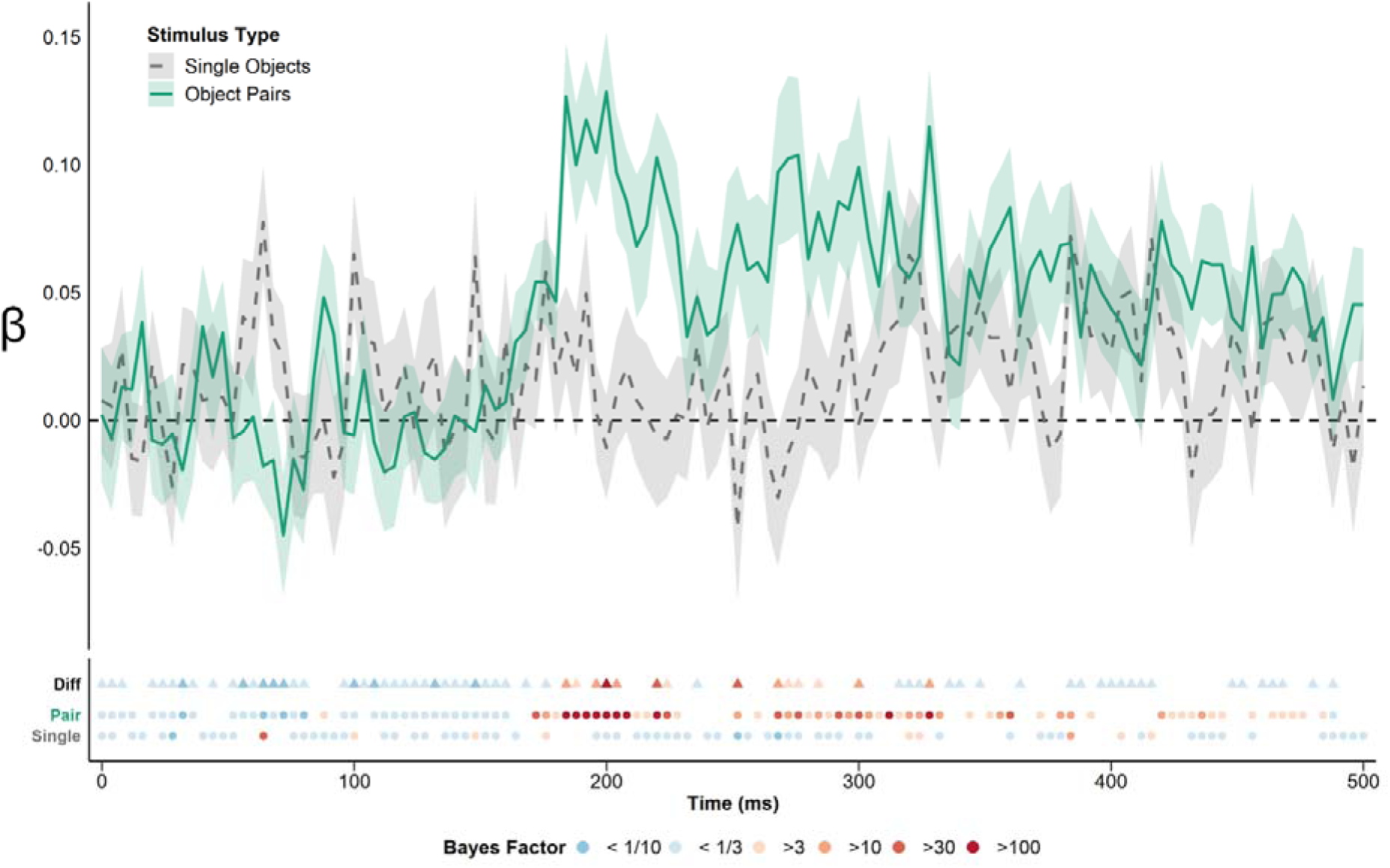
Beta values for <u>Real-world Size</u> included as a categorical predictor in (separate) multiple regression analyses for the single object (in grey) and object pair (in green) neural RDM series. Each regression also included three low-level visual control models as continuous predictors (see Figure 4). Note that the object pair betas here are identical to those in Figure 4 (green lines); shaded region is within-subjects standard error. Lower panel shows colour coded Bayes Factors for each timepoint: circles are one-sample Bayesian t-tests evaluating β > 0; triangles are paired Bayesian t-tests evaluating β_pair_ > β_single_.

#### Comparisons with a Deep Convolutional Neural Network

Finally, we considered how real-world size information about our object pair stimuli emerged in the layers of a deep convolutional neural network (AlexNet, Krizhevsky et al., 2012). Here we inspected the correlations between the real-world size model and RDMs corresponding to the 5 convolutional and 2 fully-connected layers of AlexNet. As can be seen in Figure 6B, real-world size was significantly correlated with the upper, fully-connected layers (FC6: rho = 0.20, *p* = 0.0079; FC7: rho = 0.24, *p* = 0.0016, FDR-corrected), but had little representational overlap with the early layers of AlexNet. Since low-level features differences between the large and small stimuli should be well-captured by the lowest DNN layers, the fact that these layers did not correlate with <u>Real-world Size</u> once again suggests that this model did not simply reflect visual differences between the large and small stimuli.

**Figure 6.**
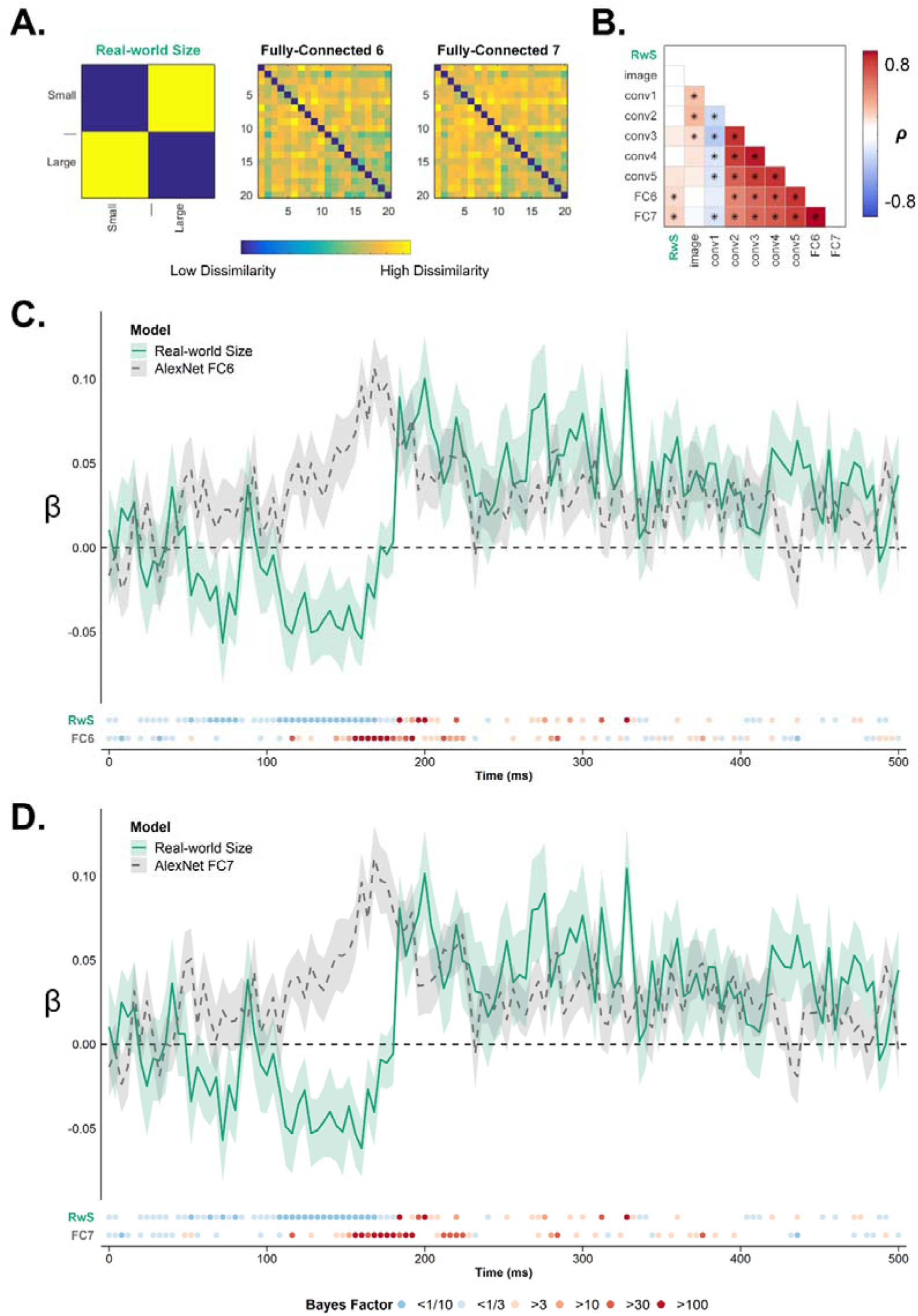
Predicting the neural response to Object Pairs using <u>Real-world Size</u> and DNN activations. A) Model RDMs based on <u>Real-world Size</u> and activations on the final two fully-connected layers of AlexNet. B) Intercorrelation matrix for <u>Real-world Size</u> and AlexNet RDMs for all layers. Asterisks indicate correlations significant at p <.01, FDR adjusted. Where <u>Real-world Size</u> was significantly correlated with both fully-connected layers, this model bore little resemblance to the lower layers of AlexNet. C&D) Beta estimates from (separate) multiple regressions of the neural RDM series for object pairs that included <u>Real-world Size</u> as a categorical predictor alongside continuous predictors based on AlexNet layers FC6 (C) and FC7 (D). Shaded region is within-subjects standard error.

To further examine the overlap of the real-world size model with AlexNet, we next ran multiple regressions of the object pair neural RDM series using <u>Real-world Size</u> as a categorical predictor alongside the later RDMs for FC6 and FC7 of AlexNet (treated as continuous predictors, see Figure 6A). Figure 6C and 6D show that in both cases, <u>Real-world Size</u> contributes unique explanatory power for the neural response evoked by object pairs, indicating that representations of spatial scale evoked by Object Pairs are not accounted for by the high-level representations that the DNN gleaned from these stimuli.

## Discussion

To apprehend a familiar object is to rapidly and automatically appreciate its inherent ‘real-world’ size (Konkle and Oliva, 2012b; Hagen et al., 2023). Investigations of both isolated objects and entire scenes have revealed real-world size to be a key organising principle of the visual system, with the spatial scale of objects – and thus their associated portability and affordances – influencing both their neural representation and the way observers recognise and respond to them (Mullally and Maguire, 2011; Konkle and Oliva, 2012a). Here we exploited the automatic retrieval of real-world size information following object recognition to examine how object processing is facilitated when the object forms part of a contextually-associated pair (i.e., an object constellation).

In Experiment 1a, we found that observers judged the real-world size of ambiguous (silhouetted) objects faster and more accurately when they appeared in contextually-associated pairs than when they appeared in isolation, suggesting that object recognition itself was more efficient when observers could integrate information across objects appearing in a familiar configuration. This inter-object contextual facilitation finding replicates that of Bar and Ullman (1996), whose participants recognised ambiguous objects better when they appeared in the context of another related object. In a follow-up experiment (1b), we verified that real-world size judgements of object pairs forming familiar object constellations (e.g., *plate + spoon*) were also more accurate than those of randomly combined objects within each size category (e.g., *plate + paintbrush*). This suggests that the object pair advantage observed in Experiment 1a did not arise simply because pair displays carry twice as much information as single object displays do, but was rather related to the presence of contextually-associated objects in a familiar configuration. Finally, our EEG experiment revealed that information about real-world size arose in the neural response to silhouetted object constellations around 170 ms after stimulus onset and peaked around 200 ms. These neural representations endured even when controlling for differences in large and small objects’ visual features, and were only evident for contextually-associated pairs of objects, and not for the individual object elements presented alone. Furthermore, around 200ms after stimulus onset, information about real-world size was significantly stronger in the neural response to contextually-associated object pairs than it was in the response evoked by isolated objects.

We interpret these results as reflecting inter-object facilitation of object recognition (Bar and Ullman, 1996). Here, integrating information across contextually-associated objects in a familiar constellation facilitated their recognition, thus making it easier to access information about the objects’ real-world size. On this interpretation, the facilitatory interaction between the processing of individual objects must have occurred shortly before 200 ms (when information about the objects’ real-world size was evident in the neural response). This is substantially earlier than facilitatory interactions between associated scenes and objects, which arise some 300 ms after stimulus onset (Brandman and Peelen, 2017, 2023). An explanation for this difference could relate to the finding that scenes and objects are processed in parallel ventral stream pathways (Levy et al., 2001; Park et al., 2011), such that interactions between scenes and objects require interactions across pathways. By contrast, interactions between objects could occur within the object processing pathway itself.

An alternative explanation worth considering is that our results reflect the accumulation of evidence (about object identity, and thus also real-world size) for both objects, but *without* facilitatory inter-object interactions. For example, each object may provide weak evidence for either large or small real-world size, with this evidence accumulating for multiple objects presented together. The data here would suggest that such evidence accumulation is non-linear; for example, evidence may have been more likely to reach a critical real-world size recognition threshold (and associated neural representation) for two objects versus one object. This parallel evidence accumulation account is undermined, however, by our findings in the behavioural follow-up experiment, in which observers’ ability to judge objects’ real-world size was significantly better for pairs forming familiar object-constellations than for pairs comprising random recombinations of objects within each size condition (i.e., “shuffled” across pairs; Kaiser et al., 2014).

We also manipulated objects’ relative depth, with the proportional retinal size of the two objects within the pair suggesting they were either at the same or different depth. We hypothesized that objects inferred to be at different depths to the viewer would be harder to integrate, similar to how disrupting objects’ relative spatial position impairs recognition (Bar and Ullman, 1996) and reduces neural measures of integration (Quek and Peelen, 2020). However, there was no evidence that this manipulation influenced either overt recognition of real-world size or the neural representations of real-world size evoked by object pairs. This suggests that the visual system is tolerant to relative size when integrating information across objects. This may reflect the variability of objects’ relative sizes in daily life, where contextually-associated objects are encountered at different depths (and thus have non-proportional retinal sizes). For example, a stapler can be observed next to a piece of paper (Figure 1B), but is also frequently observed closer or farther away from this object. On the other hand, given that texture is a strong cue for distance (Gibson, 1950; Todd and Akerstrom, 1987), the silhouette stimuli used here may have impaired or reduced the degree to which observers perceived the depth of the objects. Relative-depth effects for contextually-associated objects could well be evident when texture and other internal object cues are preserved, as in photographs.

Separately from the question of inter-object facilitation, the data here also speak to the literature on how real-world size is itself encoded in the neural response to objects. Real-world size is a dimension that pertains to all stimuli along the object-scene continuum. At the level of individual objects, real-world size is an organising principle within ventral occipitotemporal cortex (Konkle and Oliva, 2012a; He et al., 2013; Konkle and Caramazza, 2013). When retinal size is fixed, lateral ventral temporal regions respond more strongly to objects of small real-world size (e.g., a tennis ball) compared to large (e.g., a desk), while medial temporal regions exhibit the opposite activation pattern (Konkle and Oliva, 2012a). Such effects appear to reflect regional sensitivity to both featural and functional characteristics that vary systematically between small and large objects (Mullally and Maguire, 2011; Konkle and Oliva, 2012a; Troiani et al., 2014; Bainbridge and Oliva, 2015; Julian et al., 2016; Long et al., 2018; Wang et al., 2022). Studies investigating the timecourse of real-world size using single objects have shown that information about real-world size is reflected in the neural response just 120 ms after stimulus onset (e.g., Khaligh-Razavi et al., 2018; Wang et al., 2022). Such early representations likely reflect visual feature differences between large and small objects, such as degree of rectilinearity (Long et al., 2018; Wang et al., 2022). In our case, we found no evidence that neural responses evoked by isolated object silhouettes contained information about their real-world size beyond what could be accounted for by visual feature differences. This almost certainly relates to the ‘degraded’ nature of our stimuli, whose silhouette composition significantly impedes object recognition.

To our knowledge, we are the first to consider the timecourse over which information about real-world size arises for *object constellations*, showing that this property is encoded in the neural response to object pairs at around 200 ms after stimulus onset. Crucially, these results are unlikely to reflect low-or mid-level visual feature differences between large and small constellations, for several reasons. First, we selected our stimuli to avoid obvious differences between large and small conditions and used silhouettes to remove texture and colour features. Second, we included the output of multiple computational models in the EEG analyses to show that visual feature differences could not account for the real-world size representation at 200 ms. Finally, the absence of information about real-world size in the neural response to the same objects presented in isolation further argues against a feature driven effect.

If not the representation of size-covarying visual features, what drives the different neural activity patterns for large and small constellations? One possibility is that recognizing the constellations activated corresponding scene representations, such that the activity patterns reflect scene size (e.g., desktop *vs.* living room). Neural responses to scene images capture information about spatial scale as early as 100 ms after stimulus onset (Cichy et al., 2017), though as for objects, such early representations are likely driven by differences in low-level features across small and large scenes (Stansbury et al., 2013). More abstract signals of scene size (i.e., representations that generalise across image-level changes in clutter, luminance, and contrast) appear to arise later, from around 250 ms (Cichy et al., 2017), and are evident even for auditory stimuli that capture the spatial extent of real-world spaces (Teng et al., 2017). However, as noted earlier, the neural separability of small and large real-world sized objects cannot be interpreted as a direct encoding of objects’ abstract size – since there are also inherent conceptual and functional distinctions between these object classes. In the case of our stimuli, these include categorical differences (small objects mostly comprise tools/manmade artefacts; large objects are mostly furniture, Magri et al., 2021; Almeida et al., 2023), action differences (objects within small pairs can often act on each other; objects within large pairs generally do not, Baeck et al., 2013), and affordance differences (large objects were non-manipulable/had navigational affordances, Bonner and Epstein, 2017; small objects were manipulable/had grasp affordances, Haddad et al., 2024).

More generally, our results raise the question of how object and scene representations are related. By moving from single objects to object constellations, it becomes clear that the object-scene distinction is not so clear-cut. Indeed, even the processing of single objects may involve processing their real-world context – i.e., surrounding scene. Seeing a large object necessarily implies a large space, activating scene-selective cortex (Mullally and Maguire, 2011). Furthermore, object representations in scene-selective regions reflect the context within which objects are usually observed, including their co-occurrences with other objects (Aminoff et al., 2007; Bonner and Epstein, 2021; Fu et al., 2022). Our results align with these findings, showing that multiple objects viewed together activate the objects’ shared context within 200 ms.

To conclude, our results reveal the timecourse of object processing for objects comprising familiar constellations, showing that neural responses code for the objects’ real-world size around 200 ms after stimulus onset only when the objects appear in pairs, and not when they appear in isolation. We interpret these findings as reflecting inter-object facilitation between contextually-associated objects. Our study raises several new questions, including about the relationship between representations along the continuum from objects to scenes.

## Conflict of interest statement

The authors declare no competing financial interests.

## Acknowledgements

This project has received funding from the European Research Council (ERC) under the European Union’s Horizon 2020 research and innovation programme (grant agreement No. 725970). The authors thank Angeliki Karaiskou and Manon Maarseveen for their assistance with data collection, and Dr Sushrut Thorat for assisting with the layerwise AlexNet activations for our stimuli.

## Notes

https://osf.io/u4cde/

### Competing Interest Statement

The authors have declared no competing interest.

### Summary of Updates

Data from an additional control experiment incorporated.

